# Species packing and the latitudinal gradient in beta-diversity

**DOI:** 10.1101/2020.07.14.200006

**Authors:** Ke Cao, Richard Condit, Xiangcheng Mi, Lei Chen, Haibao Ren, Wubing Xu, David F. R. P. Burslem, Chunrong Cai, Min Cao, Li-Wan Chang, Chengjin Chu, Fuxin Cui, Hu Du, Sisira Ediriweera, C.S.V. Gunatilleke, I.U.A.N. Gunatilleke, Zhanqing Hao, Guangze Jin, Jinbo Li, Buhang Li, Yide Li, Yankun Liu, Hongwei Ni, Michael J. O’Brien, Xiujuan Qiao, Guochun Shen, Songyan Tian, Xihua Wang, Han Xu, Yaozhan Xu, Libing Yang, Sandra L. Yap, Juyu Lian, Wanhui Ye, Mingjian Yu, Sheng-Hsin Su, Chia-Hao Chang-Yang, Yili Guo, Xiankun Li, Fuping Zeng, Daoguang Zhu, Li Zhu, I-Fang Sun, Keping Ma, Jens-Christian Svenning

**Affiliations:** State Key Laboratory of Vegetation and Environmental Change, Institute of Botany, Chinese Academy of Sciences, Beijing 100093; Key Laboratory of Biodiversity Sciences and Ecological Engineering, Ministry of Education, College of Life Sciences, Beijing Normal University, Beijing 100875; Morton Arboretum, 4100 Illinois Rte. 53, Lisle, IL 60532, USA; Field Museum of Natural History, 1400 S. Lake Shore Dr., Chicago, IL 60605, USA; Center for Biodiversity Dynamics in a Changing World (BIOCHANGE) & Section for Ecoinformatics and Biodiversity, Department of Biology, Aarhus University, Ny Munkegade 114, DK-8000 Aarhus C, Denmark; School of Biological Sciences, University of Aberdeen, Cruickshank Building, St Machar Drive, Aberdeen AB24 3UU, UK; Institue of Natural Resources and Ecology, Heilongjiang Academy of Sciences, Harbin 150040; CAS Key Laboratory of Tropical Forest Ecology, Xishuangbanna Tropical Botanical Garden, Chinese Academy of Sciences, Kunming 650223; Center of Conservation Biology, Core Botanical Gardens, Chinese Academy of Sciences, Wuhan 430074, China; Taiwan Forestry Research Institute, 53 Nanhai Road, Taipei 100051; Sun Yat-sen University, Guangzhou 510275; Institute of Subtropical Agriculture, Chinese Academy of Sciences, Changsha, Hunan 410125; Faculty of Applied Sciences, Uva Wellassa University, Badulla 90000, Sri Lanka; Department of Botany, University of Peradeniya, Peradeniya 20400, Sri Lanka; School of Ecology and Environment, Northwestern Polytechnical University, Xi’an 710072; Center for Ecological Research, Northeast Forestry University, Harbin 150040; Research Institute of Tropical Forestry, Chinese Academy of Forestry, Guangzhou 510520; Heilongjiang Key Laboratory of Forest Ecology and Forestry Ecological Engineering, Heilongjiang Forestry Engineering and Environment Institute, Harbin 150040; Heilongjiang Academy of Forestry, Harbin 150081; Área de Biodiversidad y Conservación, Universidad Rey Juan Carlos, c/ Tulipán s/n., E-28933 Móstoles, Spain; Key Laboratory of Aquatic Botany and Watershed Ecology, Wuhan Botanical Garden, Chinese Academy of Sciences, 430074; East China Normal University, Shanghai 200241; Institute of Biology, University of the Philippines, Diliman, Quezon City, PH 1101, Philippines; South China Botanical Garden, Chinese Academy of Sciences, Guangzhou 510650; College of Life Sciences, Zhejiang University, Hangzhou 310058; National Sun Yat-sen University, Kaohsiung, 80424; Guangxi Key Laboratory of Plant Conservation and Restoration Ecology in Karst Terrain, Guangxi Institute of Botany, Guangxi Zhuang Autonomous Region and Chinese Academy of Sciences, Guilin, 541006; Department of Natural Resources and Environmental Studies, National Dong Hwa University, Hualian 97401

**Keywords:** Beta-diversity, gamma-diversity, sampling bias, latitude, species packing, niche specialization

## Abstract

The decline in species richness at higher latitudes is among the most fundamental patterns in ecology. Whether changes in species composition across space (beta-diversity) contribute to this gradient of overall species richness (gamma-diversity) remains hotly debated. Previous studies that failed to resolve the issue suffered from a well-known tendency for small samples in areas with high gamma-diversity to have inflated measures of beta-diversity. Here, we provide here a novel analytical test, using beta-diversity metrics that correct the gamma-diversity and sampling biases, to compare beta-diversity and species packing across a latitudinal gradient in tree species richness of 21 large forest plots along a large environmental gradient in East Asia. We demonstrate that after accounting for topography and correcting the gamma-diversity bias, tropical forests still have higher beta-diversity than temperate analogs. This suggests that beta-diversity contributes to the latitudinal species richness gradient as a component of gamma-diversity. Moreover, both niche specialization and niche marginality (a measure of niche spacing along an environmental gradient) also increase towards the equator, after controlling for the effect of topographic heterogeneity. This supports the joint importance of tighter species packing and larger niche space in tropical forests while also demonstrating the importance of local processes in controlling beta-diversity.

## Introduction

Beta-diversity is the variation of species composition across space, and it is a key element of conservation planning because it indicates whether diversity is concentrated within a few sites or spread across many sites (Koleff *et al.* 2003; Anderson *et al.* 2011; Socolar *et al.* 2016). One factor enhancing beta-diversity should be large niche space, i.e., more species sharing more available niches, perhaps associated with abiotic habitat heterogeneity (MacArthur 1972; Brown *et al.* 2013; Brown 2014; Alahuhta *et al.* 2017; Bracewell *et al.* 2018; Pontarp *et al.* 2019; Storch and Okie 2019). Another feature elevating beta-diversity would be dense species packing, i.e., many narrow niches result from stable climate and high productivity (Janzen 1967; MacArthur 1972; Brown 2014; Bracewell *et al.* 2018; Pontarp *et al.* 2019; Storch and Okie 2019). Both stable climate and greater productivity would then lead to higher beta-diversity at low latitudes (Gaston 2000; Willig *et al.* 2003; Hillebrand 2004; Pontarp *et al.* 2019). On the other hand, if beta-diversity is driven mostly by abiotic heterogeneity, we would not expect a latitudinal gradient in beta-diversity, since the abiotic heterogeneity should not vary with latitude. These alternatives remain unresolved and studies on the causes of the latitudinal gradient in beta-diversity appears to reach opposing conclusions (Lenoir *et al.* 2010; Kraft *et al.* 2011; De Cáceres *et al.* 2012; Mori *et al.* 2013; Myers *et al.* 2013; Qian *et al.* 2013; Sreekar *et al.* 2018). Underlying the debate has been controversy about statistical biases in tools for measuring beta-diversity.

The bias in beta-diversity metrics arises from a dependence on sample size that interacts with gamma-diversity (Condit *et al.* 2005; Kraft *et al.* 2011; Tuomisto and Ruokolainen 2012; Myers and LaManna 2016), a bias that is easy to illustrate using simple measures of species overlap. Small samples rarely (if ever) capture all local species. Two small samples from two sites that have exactly the same composition will appear to differ by randomly capturing different subsets of the local communities. The fewer the species sampled, the greater this artifactual beta-diversity will appear (Condit *et al.* 2005; Tuomisto and Ruokolainen 2012). A crucial aspect of the sample size bias is the dependence on gamma-diversity it engenders, since small samples underestimate diversity more severely in species-rich sites than in species-poor sites (Condit *et al.* 2005; Kraft *et al.* 2011; Tuomisto and Ruokolainen 2012; Chao *et al.* 2014; Sreekar *et al.* 2018). This bias has led authors to develop metrics that correct beta-diversity for sample size (Condit *et al.* 2005; Chao *et al.* 2014; Cao *et al.* 2021) or tools based on comparisons with null models (Kraft *et al.* 2011; Myers and LaManna 2016). Crucial in the sample size bias is the dependence on gamma-diversity it engenders, since larger samples are needed in species-rich sites (Condit *et al.* 2005; Tuomisto and Ruokolainen 2012; Chao *et al.* 2014; Sreekar *et al.* 2018). Once correcting for sample size bias, gamma-diversity dependence should be removed, and it should be straightforward to compare beta-diversity across a gradient of species diversity in order to evaluate the importance of species packing and total niche space.

We carry out this comparison using a steep latitudinal gradient in tree species richness, as documented in our census of 3 million trees at 21 sites spanning 50° of latitude in East Asia (Anderson-Teixeira *et al.* 2015; Feng *et al.* 2016). We define beta-diversity within each plot, so it is a measure of how tree species partition local niche space, then we compare the local estimates of beta-diversity across the latitudinal gradient. In a previous simulation study, Cao et al. (Cao *et al.* 2021) identified that the corrected beta-Shannon diversity index is highly effective at removing the bias arising from beta-diversity metrics in small samples of high gamma-diversity communities (Cao *et al.* 2021). With this corrected index, we can answer two fundamental questions about variation in beta-diversity and its impact on the overall species richness: 1) Is there a latitudinal gradient in within-plot beta-diversity? 2) Do local environmental heterogeneity, niche marginality (the distance between the species optima relative to the overall mean habitat), and niche specialization contribute to the latitudinal patterns of beta-diversity? By simultaneously testing the importance of local heterogeneity and latitude, we can establish whether species packing and total niche space contributes to higher richness in tropical relative to temperate forests.

## Materials and Methods

### Forest dynamic plots

We used data from 21 forest dynamics plots (15-52 ha) that are part of the ForestGEO and Chinese Forest Biodiversity Monitoring Networks (Anderson-Teixeira *et al.* 2015; Feng *et al.* 2016) (figure 1a; electronic supplementary material, table S1). All stems with diameter at breast height (DBH) ≥ 1 cm were spatially mapped, tagged, measured and identified to species (Condit 1998). The plots range from tropical rain forest at 2.98° N latitude to boreal forest at 51.82° N latitude (electronic supplementary material, table S1), from sea level to more than 1400 m elevation, and local topographic variation is as low as 17.7 m and as high as 298.6 m (figure 1b and electronic supplementary material, table S1). We divided plots into non-overlapping quadrats of different scales (grain sizes) (10 m × 10 m, 20 m × 20 m, and 50 m × 50 m) in order to assess the effect of grain size on beta-diversity (De Cáceres *et al.* 2012; Sreekar *et al.* 2018). We define alpha diversity as the quadrat level diversity, and gamma diversity as plot level diversity. In the main results, we present only the results at grain size of 20m × 20m, and details of results at grain size of 10m × 10m and at grain size of 50m × 50 m could be found in electronic supplementary materials (Table S2, Figure S2).

**Figure 1.**
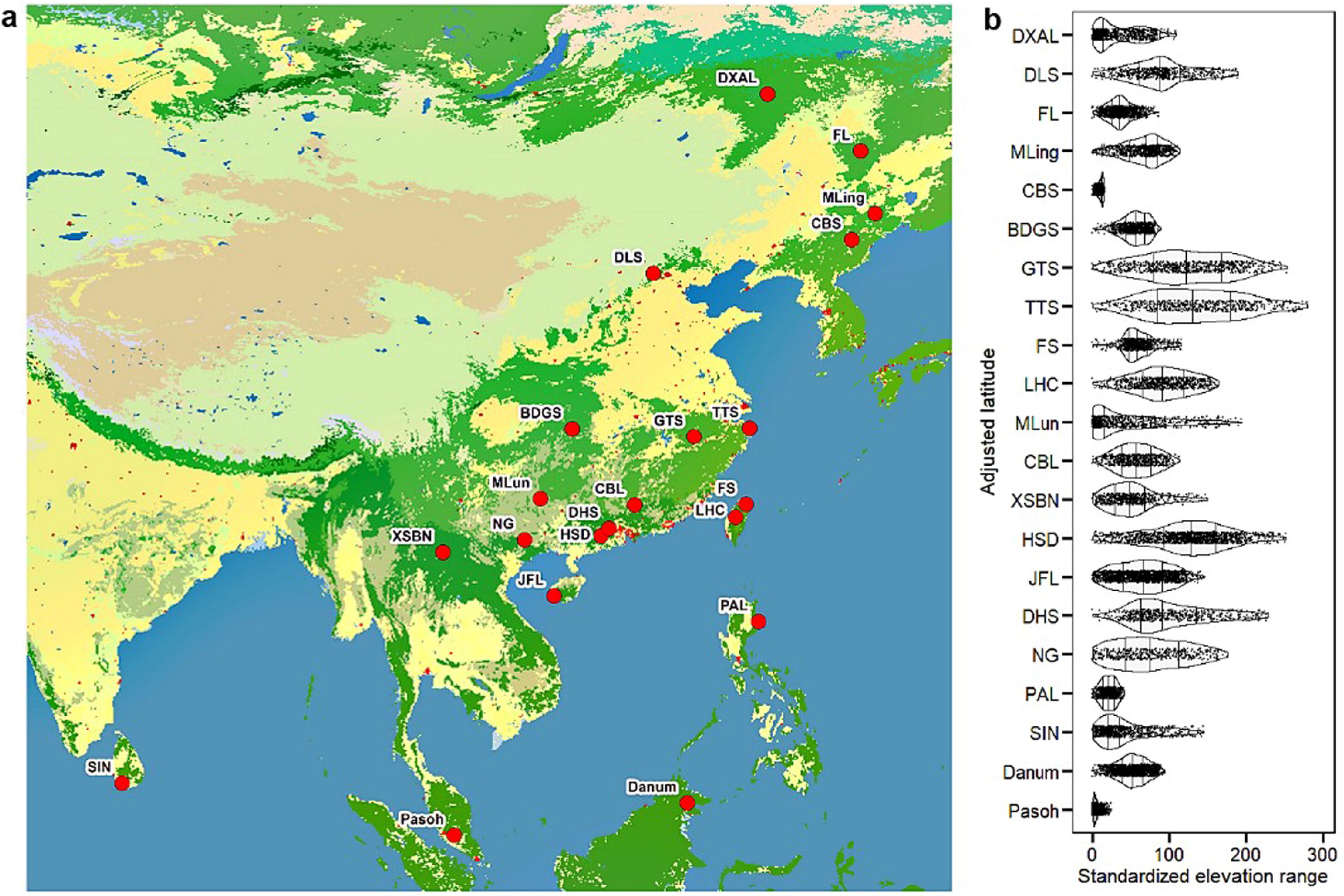
The spatial distribution of forest dynamics plots (a), and their elevational ranges (b). Panel (b) shows the latitudinal pattern of elevation range, which was transformed by subtracting the minimum elevation of each plot. The width of each violin plot reflects probability density distribution of mean elevation for 20 m × 20 m subplots in each forest dynamics plot. Full plot names are listed in electronic supplementary material, table S1.

Plot latitudes were adjusted for mean elevation: adding 100 km of latitude per 100-m increase in elevation. Local environmental heterogeneity was quantified in terms of topography, which was the only environmental factor consistently available across all plots. Specifically, we used the ratio of surface area to planimetric as a metric of topographic heterogeneity, calculating at grain sizes of 10 m × 10 m, 20 m × 20 m, and 50 m × 50 m, which provided a useful measure of the range and roughness of the overall plot, based on digital elevation models (DEMs) (Jenness 2004; Brown *et al.* 2013). Local habitat and species niches were defined using six topographic factors as environmental variables: mean elevation, convexity, slope, aspect, topographical wetness index (TWI) and altitude above channel (ACH) (Legendre *et al.* 2009; Kanagaraj *et al.* 2011; Punchi-Manage *et al.* 2013).

### Measurement of beta-diversity

To remove gamma-diversity dependence caused by sample-size bias of beta-diversity metrics, we used the correction method designed for Shannon diversity index based on the relationship between cumulative diversity and sample size (Chao *et al.* 2013). The beta-Shannon diversity index measures the heterogeneity of pooled communities, and is calculated as the effective number of compositionally distinct and equally abundant communities (Jost 2007; Tuomisto 2010):

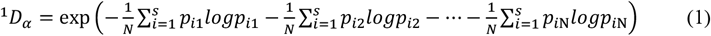

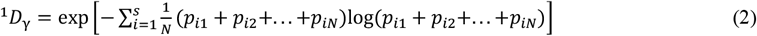

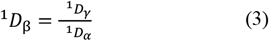

where ^1^*D*_α_, ^1^*D*_β_ and ^1^*D*_γ_ are alpha-, beta-, and gamma-Shannon diversity, respectively; *p*_i_ is the proportional abundance of species *i*; *S* and *N* are the total number of species and the total number of local communities (or plots), respectively, in the pooled communities. Alpha- and gamma-Shannon diversity are mathematically independent (i.e., gamma-diversity does not contain information of alpha-diversity) (Jost 2007). Beta-Shannon diversity weights all species by their abundance. We then used a sample-size dependence correction method to reduce the bias in beta-Shannon diversity for comparing beta-diversity among regions (Chao *et al.* 2013; Chao *et al.* 2014). As in a species accumulation curve, the expected cumulative alpha- or gamma-diversity was depicted as a function of sample size, while sample completeness was estimated from community structures of samples (Chao *et al.* 2013; Chao *et al.* 2014). Beta-diversity was then estimated from asymptotically approximated alpha- and gamma-diversity based on the diversity-sample size curve. Details of undersampling correction method for the beta-Shannon diversity can be found in the electronic supplementary material S1. Simulation work conducted by Cao et al. (2021) confirmed that β-metrics that incorporate an undersampling correction method were more effective at removing dependence on gamma-diversity and inferring casual mechanisms compared to other uncorrected beta-diversity metrics or null models (Cao *et al.* 2021).

### Community-level niche differentiation

Niche differentiation was described using attributes of specialization and marginality. Niche specialization was defined as *SD(available habitat)/SD(habitat used),* in which *SD(available habitat)* represented the standard deviation of environmental conditions for a community and *SD(habitat used)* represented the standard deviation of environmental conditions occupied by a species (illustrated in figure 2). Niche marginality was defined as the distance between a species’ optimum and the mean environmental conditions within the plot (figure 2) (Hirzel *et al.* 2002; Devictor *et al.* 2010). Both specialization and marginality were calculated from multivariate measures of habitat, known as ecological niche factor analysis (Hirzel *et al.* 2002). To better meet the assumption of normality of residual in regression model and approximate the linear relationship between niche specialization and explanatory variables (Supplementary material, figure S1a, 1c, 1e), the log- and Box-Cox transformations (Box and Cox 1964) were applied for niche specialization across grain sizes (Supplementary material, figure S1b, 1d, 1f). Based on the precise mapping of all individuals in these plots, the community-level niche marginality and specialization were respectively quantified as species-level niche marginality and specialization weighted by relative species abundance. Higher community-level niche specialization indicates the fine partitioning of available niche space, while higher community-level niche marginality indicates a larger deviation from mean environmental conditions of a community, and thus suggesting a larger niche space. Topographic variables are typically strongly correlated with the variation in resources such as water availability and soil conditions (Wright 2002; Fortunel *et al.* 2018), thus can capture potentially important axes of niche differentiation. Aspect was computed as sin(aspect) and cos(aspect), and other topographic variables were Box-Cox transformed before being included into analyses (Box and Cox 1964).

**Figure 2.**
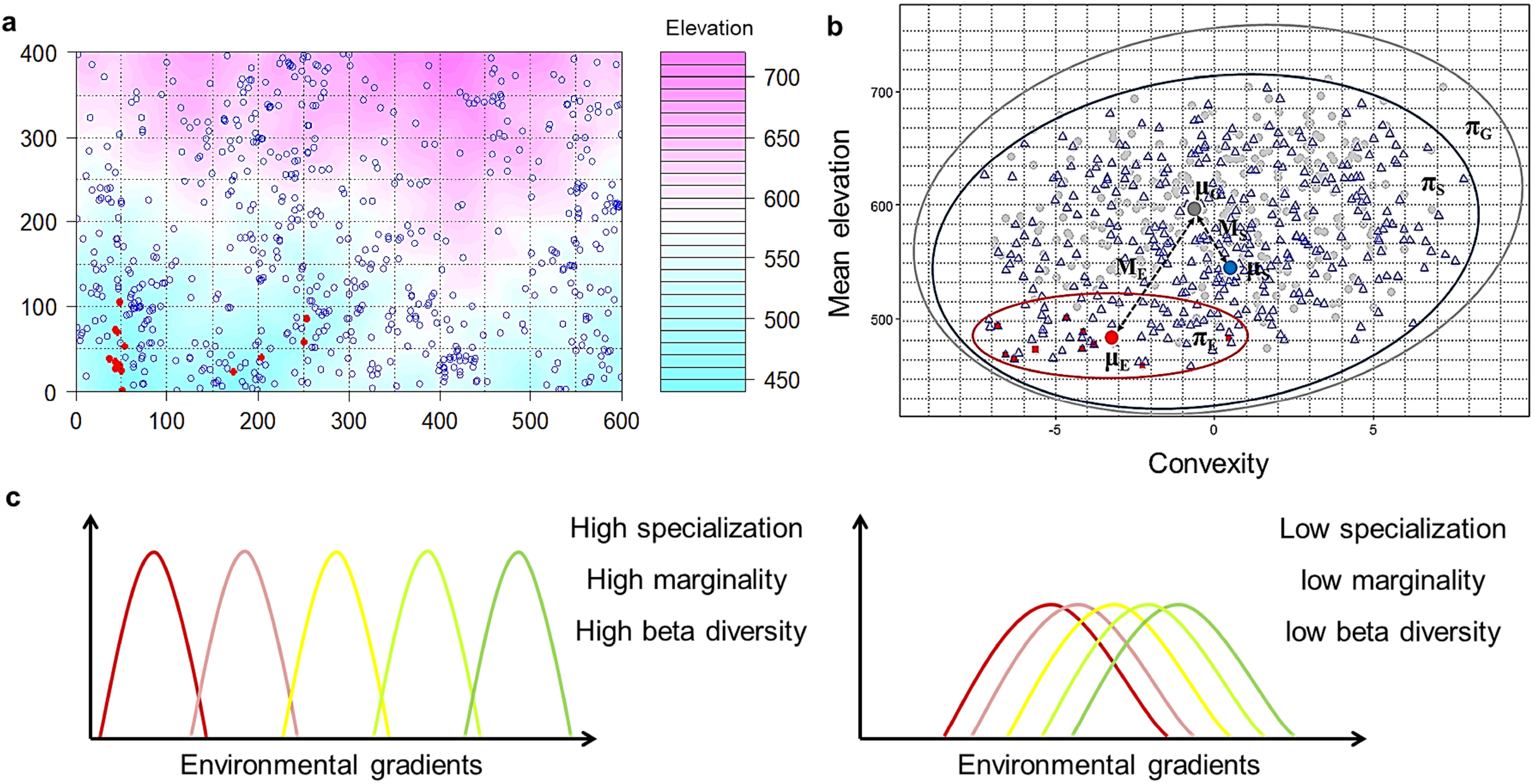
Illustration of niche specialization and marginality of *Euonymus oblongifolius* and *Symplocos stellaris* in the Gutianshan forest dynamics plot (600 m × 400 m). a. Red solid points represent the spatial distribution of *E. oblongifolius*, and blue circles represent the spatial distribution of *S. stellaris*. b. Illustration of niche specialization and marginality of *E. oblongifolius* and *S. stellaris* in two-dimensional niche space based on mean elevation and convexity of distributed 20 m × 20 m quadrats. Niche marginality is the distance from the mean habitat of the focal species to the mean habitat of community habitats. *μ_E_*, *μ_S_* and *μ_G_* represent centroids of environmental conditions for *E. oblongifolius*, *S. stellaris* and the entire community, and distances *M_E_* and *M_S_* are niche marginalities of two species. Likewise, niche specialization is ratio of the entire habitat range of a community to habitat range of the focal species. *π_E_, π_S_* and *π_G_* stand for the distributional range of for *E. oblongifolius, S. stellaris* and the entire community in two-dimensional niche space respectively, the ratio of *π_G_/π_E_* and *π_G_/π_S_* are niche specialization of two species. Grey points indicate the topographic variation of the entire community, red squares show higher niche specialization and marginality of *E. oblongifolius*, whereas blue triangles indicate lower specialization and marginality of *S. stellaris*. c. Hypothetical relationships between beta-diversity and niche. Higher community-level niche specialization indicates the fine partitioning of available niche space, while higher community-level niche marginality suggests a larger niche space. Therefore, higher specialization and marginality lead to a higher beta-diversity (left), while lower specialization and marginality lead to a lower beta-diversity (right).

#### Statistical analysis

To examine the significance of latitudinal gradients in local beta-diversity, niche specialization and niche marginality, we first modeled beta-diversity, community-level niche specialization and niche marginality against topographic heterogeneity and adjusted latitude separately using simple linear regression models. Subsequently, to determine the relative effect sizes of adjusted latitude and topography, we performed multiple linear regression models with beta-diversity, niche specialization, and niche marginality as response variables, respectively, and all variables were scaled using *(x* – *mean(x))/SD(x)* before being included.

All statistical analyses were performed with R software, version 3.6.4 (R Core Team 2019). The corrected beta-Shannon diversity was calculated using R package ‘entropart’ and ‘vegan’ (Marcon and Hérault 2015; Oksanen *et al.* 2018). The topographic variables were computed using the ‘RSAGA’ package (Brenning 2008) and the SAGA GIS software (Conrad *et al.* 2015). Ecological niche factor analysis was implemented to calculate niche metrics using R package ‘adehabitatHS’ (Calenge 2006).

### Results

Gamma-diversity declined by more than forty-fold from tropical to temperate latitudes, from 818 species at Pasoh to 18 at Daxinganling (electronic supplementary material, table S1). Beta-diversity measured by the corrected beta-Shannon diversity also declined with latitude, although this pattern was not significant (figure 3a). However, the corrected beta-Shannon diversity was significantly correlated with latitude (e.g., 20m × 20m, *standardized effect size = −0.39, p = 0.033)* in multiple regression models, after controlling for the effect of local topographic heterogeneity (electronic supplementary material, figure S2c). We also found that beta-diversity was positively correlated with community-level niche specialization, niche marginality and local topographic heterogeneity (figures 3b-3d, electronic supplementary material, figure S3). We obtained similar results across three grain sizes although the effect size of topographic heterogeneity and latitude varied with grain sizes (electronic supplementary material, Figs. S2, S3).

**Figure 3.**
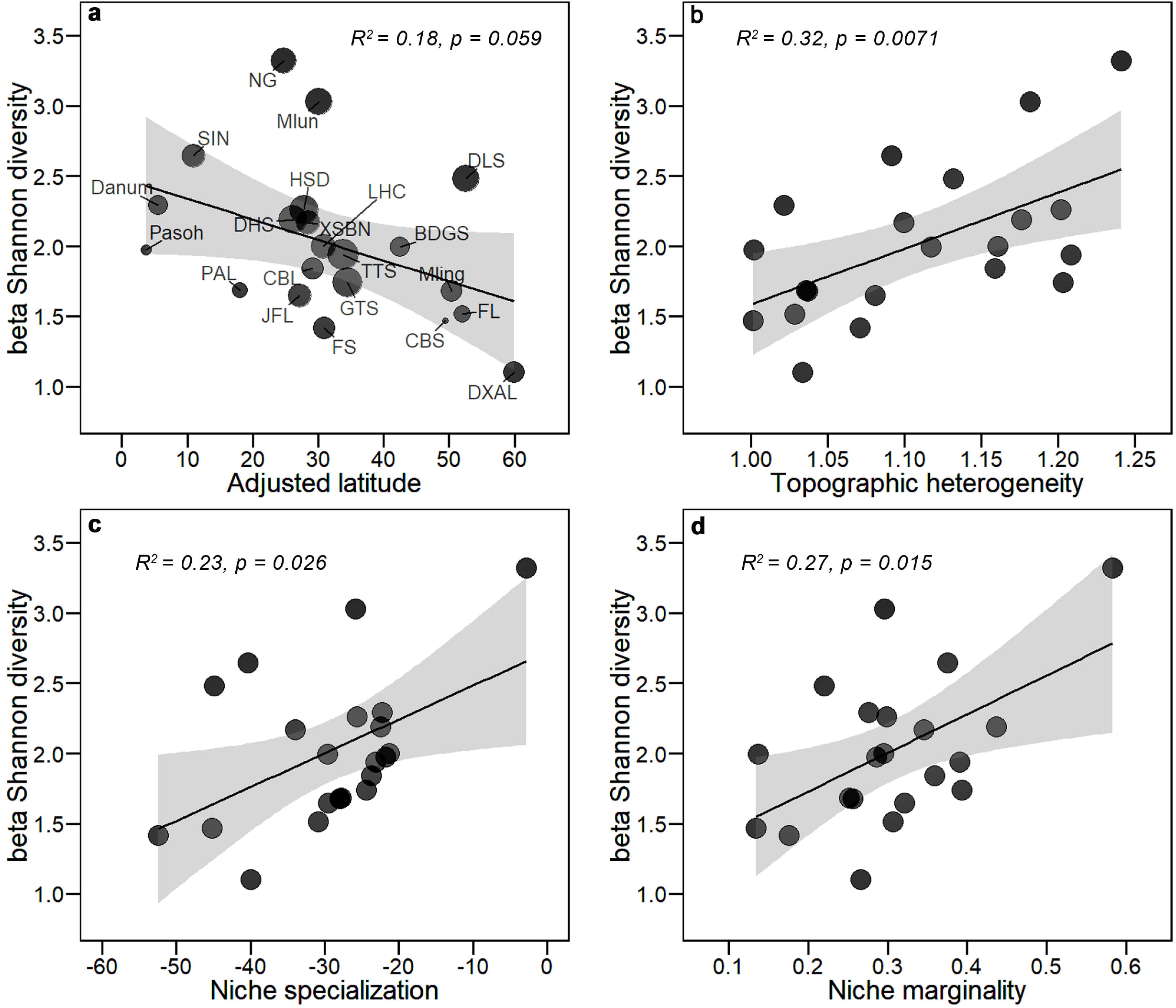
Relationships of beta-diversity (measured corrected beta-Shannon diversity) with adjusted latitude (a), and local topographic heterogeneity (b), community-level niche specialization (c), and niche marginality (d) at grain size of 20 m × 20 m. In each panel, different colours of points and lines represent grain sizes. In panels a and b, solid and dashed lines indicate significant and insignificant linear correlations (significance level, *α* = 0.05), respectively, and the shaded areas represent the 95% confidence intervals of the predictions (electronic supplementary material, table S2). Full plot names in (a) are listed in electronic supplementary material, table S1. Community-level niche specialization was Box-Cox transformed in (c).

Various predictors of beta-diversity were also associated with latitude. Both community-level niche specialization and niche marginality significantly decreased from tropical to temperate forests at some grain sizes (figures 4a, 4c, S6a, S6c). However, topographic heterogeneity did not have a significant relationship with latitude (electronic supplementary material, figure S5). Both niche specialization and niche marginality were positively correlated with each other (electronic supplementary material, figures. S4g-4i), and both were also positively associated with local topographic heterogeneity (figures 4b, 4d, electronic supplementary material, figures. S4a-4f). Multiple linear regression models confirmed these results, showing that specialization and marginality both significantly declined with latitude after controlling for topographic heterogeneity at most grain sizes. In the multiple regression models, the effect sizes of topographic heterogeneity were larger than those of adjusted latitude in predicting specialization and marginality (electronic supplementary material, Table S3, figures S6b and 6d).

**Figure 4.**
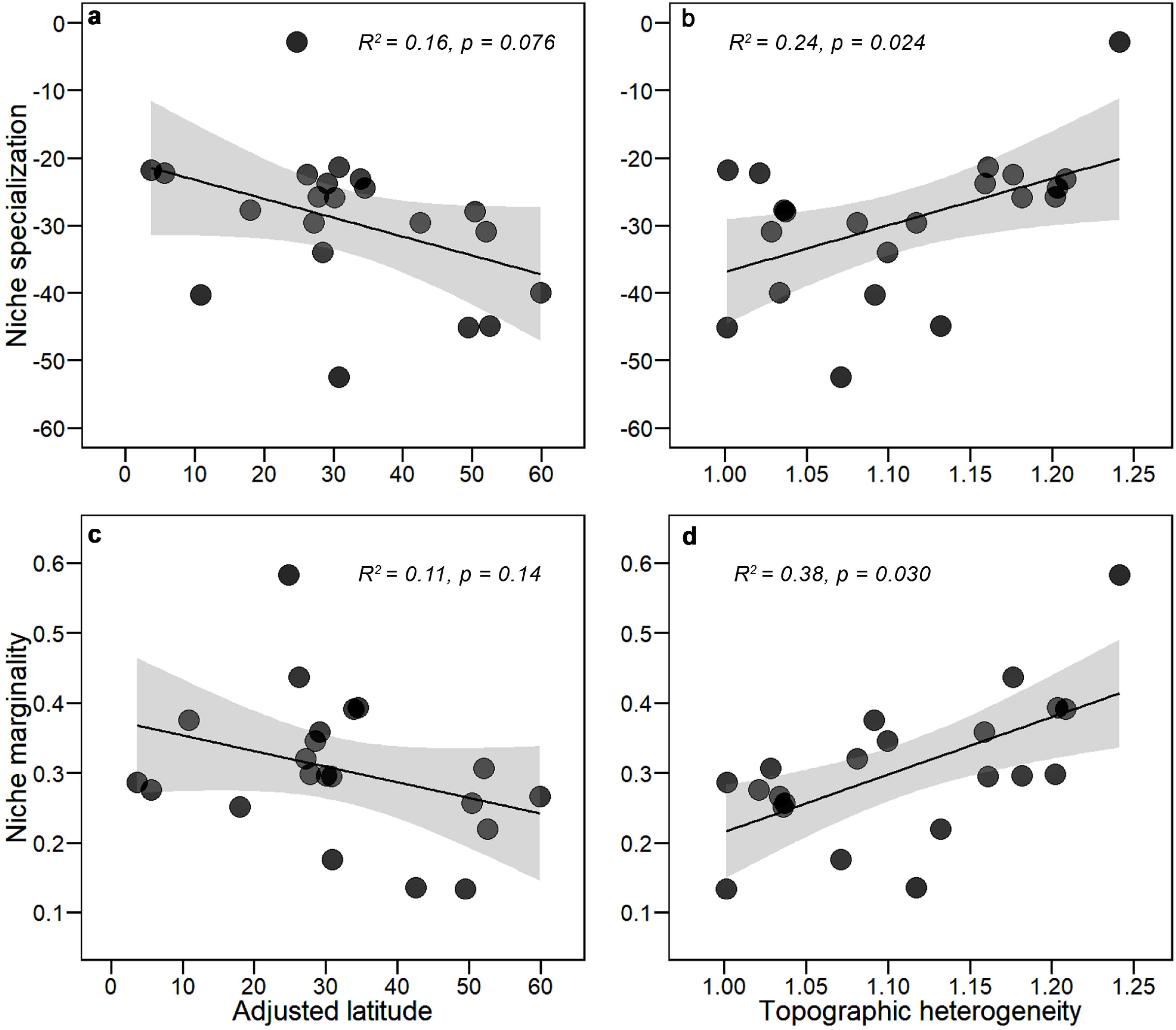
The relationships of community-level niche specialization (a and b) and marginality (c and d) with adjusted latitude and local topographic heterogeneity at grain size of 20 m × 20 m. Community-level niche specialization was Box-Cox transformed. In each panel, *R^2^* and *p-value* of the linear regression models was shown at each panel, and shaded areas represent the 95% confidence intervals of the predictions (electronic supplementary material, table S4).

## Discussion

Whether beta-diversity contributes to the latitudinal diversity gradient has been intensely debated in recent years, largely because of the bias in beta-diversity metrics in small samples of high gamma-diversity communities (Condit *et al.* 2005; Kraft *et al.* 2011; Tuomisto and Ruokolainen 2012; Qian *et al.* 2013; Myers and LaManna 2016; Sreekar *et al.* 2018). To move this debate forward, we first examined the latitudinal gradient in beta-diversity by removing the gamma-diversity and sample size bias with a correction for undersampling (Chao *et al.* 2013; Chao *et al.* 2014), while also accounting for the effect of topographic heterogeneity statistically. Our results showed that beta-diversity increased from high to lower latitudes, in lines with a number of previous studies also finding higher beta-diversity in the tropics (Koleff *et al.* 2003; Willig *et al.* 2003; Vazquez and Stevens 2004; Myers *et al.* 2013). This supports the hypothesis that beta-diversity contributes to the latitudinal gradient in species richness. Since topographic heterogeneity did not systematically vary with latitude, it appears that local topographic heterogeneity does not contribute to the latitudinal gradient in beta-diversity, in line with previous findings (Alstad *et al.* 2016; Chu *et al.* 2019).

High beta diversity in the tropics reveals higher species turnover at lower latitudes, meaning tighter species packing and expanded niche space in tropical relative to temperate forests (Ricklefs and Schluter 1993; Gaston 2000; Vazquez and Stevens 2004; Brown 2014; Pontarp *et al.* 2019). These hypotheses have been investigated for decades, with dense species-packing in large niche space attributed to stable climate and higher productivity in the tropics (MacArthur 1965; Ricklefs and Schluter 1993; Evans *et al.* 2005; Brown 2014; Pontarp *et al.* 2019). We found increasing niche marginality and specialization towards lower latitudes, supporting this hypothesis. Perhaps larger niche space enables more species to utilize more variable resources, while higher niche specialization allows species to specialize on narrower subsets of the resources available (MacArthur 1965; Ricklefs and Schluter 1993; Evans *et al.* 2005; Brown 2014; Pontarp *et al.* 2019). These consequently reduce niche overlap and competition between co-occurring species and facilitates species coexistence (Arellano *et al.* 2017). Tighter species packing and larger niche space in the tropics could be related to other mechanisms as well, such as higher diversification rate (Fine 2015) and stronger conspecific negative density dependence (Fine *et al.* 2004; Umaña *et al.* 2017) at lower latitudes.

We also conclude that beta-diversity at extent of 15-52 ha is largely driven largely by local processes—specifically, topographic heterogeneity and the niche differentiation it fosters. However, topographic heterogeneity did not contribute to the latitudinal gradient in beta-diversity (figures 3 and 4). This may seem an unsurprising result, but the roles of local ecological processes have been questioned given the broad latitudinal gradient of gamma-diversity (Gaston 2000; Kraft *et al.* 2011). We suggest that the effect of local processes have been obscured by the biases in beta-diversity metrics of small samples from high gamma-diversity communities in previous studies (Myers and LaManna 2016). Moreover, our large samples over 55 degrees of latitude provide comparable measures of niche differentiation, topographic heterogeneity, and beta-diversity, well beyond what was available in early studies (Brown *et al.* 2013; Shen *et al.* 2013). Our results could be refined by considering the influence of additional factors that contribute to local environmental heterogeneity and niche differentiation, such as soil types and soil nutrients (Baldeck *et al.* 2013), which could also contribute to beta-diversity. The biases in beta-diversity metrics in small sample from high gamma-diversity communties are also associated with other attributes of communities such as the species abundance distributions (Chao and Jost 2012), and tests of the alternative techniques in other systems are warranted.

In conclusion, our results support that a latitudinal gradient in beta-diversity contributes to the latitudinal gradient in tree species richness after separately controlling for local topographic heterogeneity and the bias in beta-diversity metrics in small samples of high gamma-diversity areas. Our results further suggest tighter species packing and larger niche space in tropical forests (MacArthur 1965; Ricklefs and Schluter 1993; Gaston 2000), but also confirmed environmental heterogeneity as a determinant of beta-diversity. Our findings help resolve the ongoing debates on the contribution of local beta-diversity to latitudinal gradient of species richness.

## Data availability

The data supporting Figure 1-4 and code for data analyses are available from the Dryad Digital Repository: https://doi.org/10.5061/dryad.tht76hdww. Full census data are available upon reasonable request from the data portal of ForestGEO (http://ctfs.si.edu/datarequest/) and CForBio (Chinese Forest Biodiversity Monitoring Networks (http://www.cfbiodiv.org).

## Author contributions

K.C., R.C., X.M., K.M. and J.C.S. designed research, K.C. and X.M. compiled and analysed data; K.C., R.C., X.M., K.M. and J.C.S. wrote the draft with substantial input from L.C., W.X., D.F.R.P.B. and M.J.B.. Many authors contributed to data collection of forest censuses and all authors contributed to revisions of the manuscript.

## Acknowledgements

We thank Dingliang Xing, Tak Fung, Fangliang He and Gabriel Arellano for comments on earlier draft. We thank Alex Karolus for leading the census in Danum Valley forest plot, and we are grateful to Mike Bernados and Bill McDonald for species identifications, to Fangliang He, Stuart Davies and Shameema Esufali for advice and training, to Qianjiangyuan National Park, Fushan Research Center, Lienhuachih Research Center and Sri Lankan Forest Department for logistical support and the hundreds of field-workers and students who measured and mapped the trees analyzed in this study.

