## Supplemental file for "Species packing and the latitudinal gradient in beta-diversity"

### Electronic supplementary material

### S1. Undersampling correction methods for β-Shannon diversity

Species accumulation curves have been applied to rarefy and extrapolate species richness with respect to sample size [1]. Chao *et al*. [2, 3] extended the species accumulation curve to the diversity accumulation curve, which corrects for the undersampling bias by asymptotically estimating the real α- and γ-Shannon diversity ($^{1}\hat{D}(\infty)$) of samples in a region. The $^{1}\hat{D}(\infty)$ ($q$=1, $q$ is the diversity order) is calculated as:

$$^{1}\hat{D}(\infty) = \exp[\hat{H}(\infty)] \qquad (1)$$

$\hat{H}(\infty)$ is a nearly unbiased estimator of Shannon entropy [2]:

$$\hat{H}(\infty) = \sum_{k=1}^{n-1}\frac{1}{k}\sum_{1\le X_i\le n-1}\frac{X_i}{n}\frac{\binom{n-X_i}{k}}{\binom{n-1}{k}} + \frac{f_1}{n}(1-A)^{-n+1}\left\{-\log(A) - \sum_{r=1}^{n-1}\frac{1}{r}(1-A)^r\right\} \qquad (2)$$

where $X_i$ is the species frequency of species *i*, *k* is the size of a random sample from the observed community, $f_1$ is the number of singletons (i.e., species represented by only one individual in the observed sample), and $f_2$ is the number of doubletons (i.e., species represented by only two individuals in the observed sample). $A$ is the estimated mean relative frequency of the singletons in the sample:

$$A = 2f_2/[(n-1)f_1 + 2f_2\ ] \qquad (3)$$

**Supplementary Table S1:** Basic information of 21 forest dynamic plots.

| Plot name | Area (ha) | Latitude (°N) | Longitude (°E) | Mean elevation (m) | Elevation range (m) | Gamma-diversity | Number of stems |
|---|---|---|---|---|---|---|---|
| Pasoh | 50 | 2.98 | 102.31 | 80 | 24 | 818 | 335400 |
| Danum Valley | 50 | 5.1 | 117.69 | 54.1 | 101.12 | 642 | 234916 |
| Sinharaja | 25 | 6.4 | 80.4 | 499.5 | 151 | 239 | 250131 |
| Palanan | 16 | 17.04 | 122.38 | 111 | 55 | 415 | 66000 |
| Jianfengling | 60 | 18.73 | 108.9 | 932 | 150.4 | 290 | 439676 |
| Xishuangbanna | 20 | 21.6 | 101.57 | 765.1 | 159.87 | 467 | 95834 |
| Nonggang | 15 | 22.42 | 106.95 | 260 | 190 | 223 | 67870 |
| Heishiding | 50 | 22.7 | 111.99 | 568.8 | 263 | 236 | 264391 |
| Dinghushan | 20 | 23.17 | 112.52 | 339 | 240 | 195 | 71617 |
| Lienhuachih | 25 | 23.91 | 120.88 | 765.4 | 178 | 144 | 153268 |
| Chebaling | 20 | 24.72 | 114.22 | 488 | 131 | 222 | 86517 |
| Fushan | 25 | 24.76 | 121.56 | 675.3 | 133 | 110 | 114500 |
| Mulun | 25 | 25.13 | 108 | 547 | 208.8 | 254 | 144679 |
| Gutianshan | 24 | 29.25 | 118.12 | 580.6 | 268.6 | 159 | 140700 |
| Badagongshan | 25 | 29.77 | 110.09 | 1414 | 101 | 241 | 186556 |
| Tiantongshan | 20 | 29.81 | 121.79 | 447.25 | 298.63 | 152 | 115536 |
| Donglingshan | 20 | 40 | 115.43 | 1395 | 219.3 | 53 | 52136 |
| Changbaishan | 25 | 42.22 | 128.53 | 801.5 | 17.7 | 52 | 38902 |
| Muling | 25 | 43.95 | 130.07 | 719.5 | 123 | 57 | 63877 |
| Fenglin | 30 | 48.08 | 129.12 | 439 | 66 | 46 | 94920 |
| Daxinganling | 25 | 51.82 | 122.98 | 896.7 | 115.3 | 18 | 126532 |

**Supplementary Table S2:** The results of simple linear regression models for beta-diversity across grain sizes in figure S2a-b.

| Explanatory variables | Grain size | Coefficients | Standard error | t-value | p-value | R-square |
|---|---|---|---|---|---|---|
| **Adjusted latitude** | 10m × 10m | -0.024 | 0.01 | -2.59 | 0.02 | 0.26 |
| | 20m × 20m | -0.015 | 0.01 | -2.01 | 0.06 | 0.18 |
| | 50m × 50m | -0.0087 | 0.01 | -1.86 | 0.08 | 0.15 |
| **Topographic heterogeneity** | 10m × 10m | 3.62 | 1.65 | 2.19 | 0.04 | 0.20 |
| | 20m × 20m | 4.00 | 1.33 | 3.02 | 0.007 | 0.32 |
| | 50m × 50m | 4.63 | 1.08 | 4.28 | <0.001 | 0.49 |

**Supplementary Table S3:** The results of simple regression models for niche specialization and marginality against adjusted latitude in Figure S6a and S6c.

| Response variables | Grain size | Coefficients | Standard error | t-value | p-value | R-square |
|---|---|---|---|---|---|---|
| **Niche specialization** | 10m × 10m | -0.0057 | 0.0027 | -2.10 | 0.049 | 0.19 |
| | 20m × 20m | -0.0056 | 0.0030 | -1.88 | 0.076 | 0.16 |
| | 50m × 50m | -0.0065 | 0.0027 | -2.38 | 0.028 | 0.23 |
| **Niche marginality** | 10m × 10m | -0.0033 | 0.0015 | -2.15 | 0.044 | 0.20 |
| | 20m × 20m | -0.0022 | 0.0014 | -1.55 | 0.14 | 0.11 |
| | 50m × 50m | -0.0031 | 0.0015 | -2.04 | 0.055 | 0.055 |

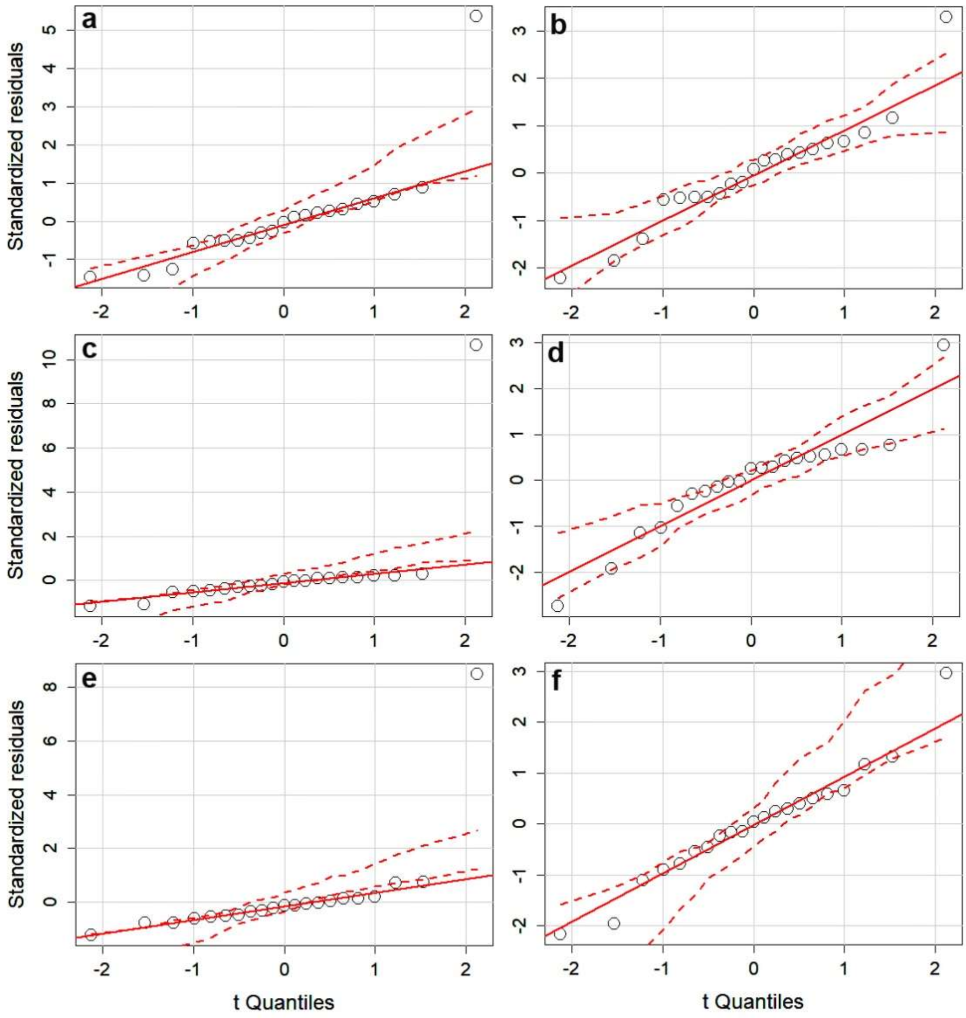


**Supplementary Figure S1**. **QQ-plot of residuals from the linear models (niche specialization ~ latitude) before (a, c, e) and after (b, d, f) Box-Cox transformation of niche specialization at grain size 10m × 10m (a, b), 20m × 20m (c, d) and 50 m × 50 m (e, f).**

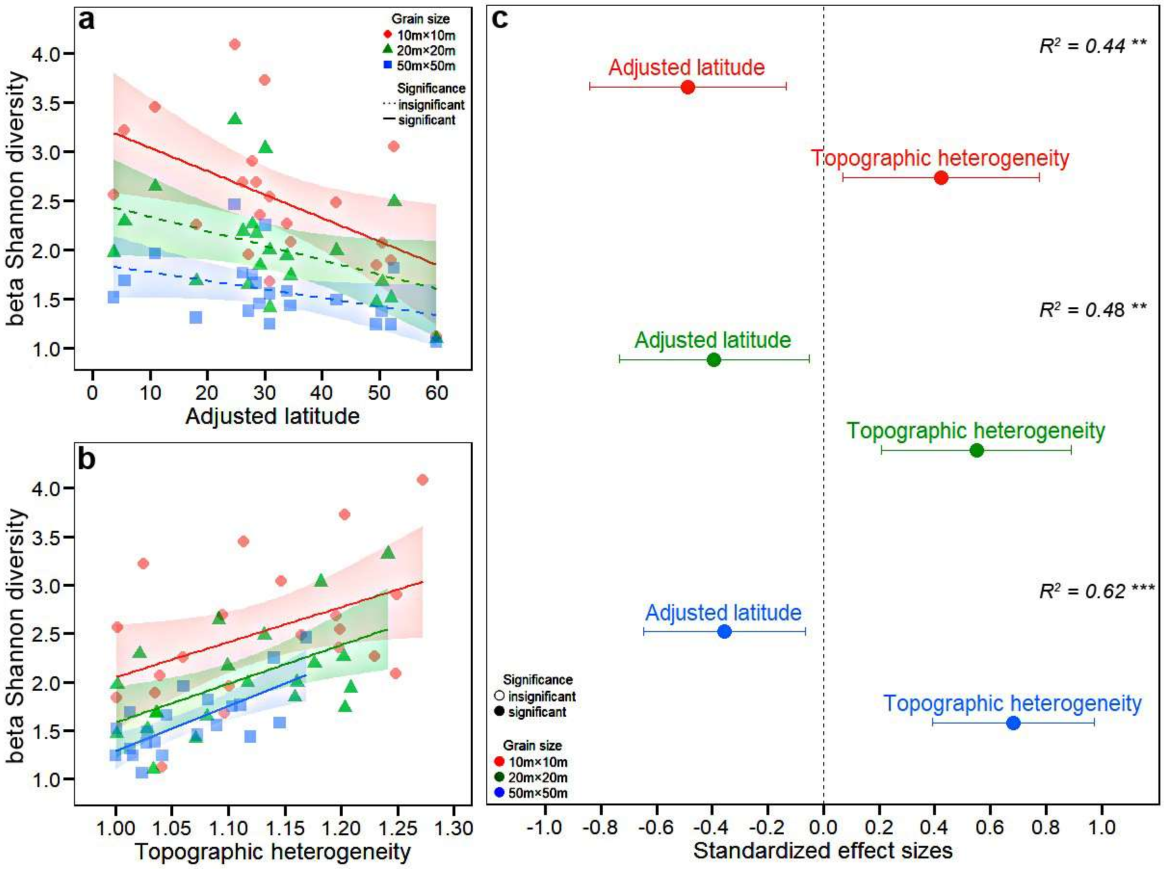


**Supplementary Figure S2. Relationship of beta-diversity (measured corrected beta-Shannon diversity) and adjusted latitude (a), and local topographic heterogeneity (b) and their effect sizes (c) across grain sizes.** In each panel, different colours of points and lines represent grain sizes. In panels a and b, solid and dashed lines indicate significant and insignificant linear correlations (significance level, $\alpha = 0.05$), respectively, and the shaded areas represent the 95% confidence intervals of the predictions (electronic supplementary material, table S2). In panel c, points represent the standardized effect sizes of explanatory variables that are significantly (solid circles) and non-significantly (open circles) different from zero, respectively. The significance level of the total $R^2$ are $\alpha < 0.001$, '***'; $\alpha < 0.01$, '**'; $\alpha < 0.05$ '*'.

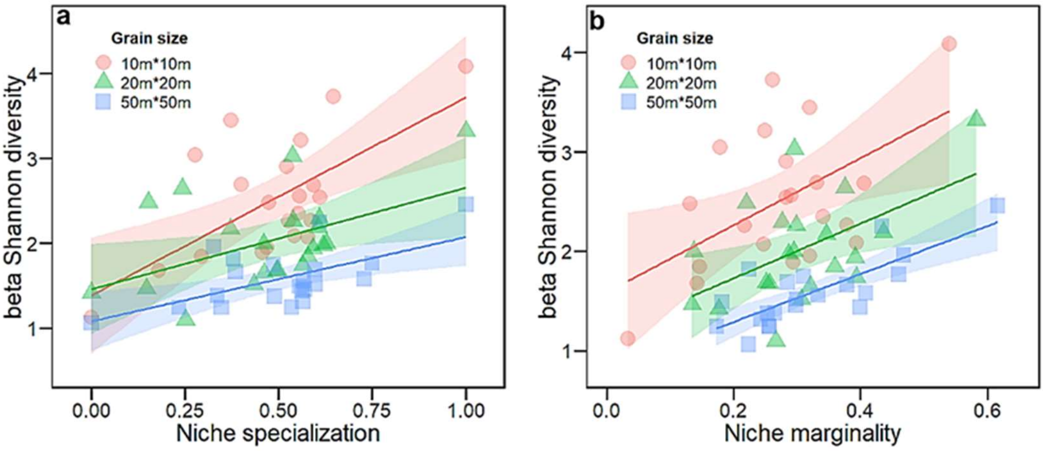


**Supplementary Figure S3. The relationship of beta-diversity (measured by the corrected beta-Shannon diversity) with (a) community-level niche specialization, and (b) community-level niche marginality across grain sizes.** In each panel, different colours of points and lines represent grain sizes. In panels a and b, solid and dashed lines indicate significant and non-significant linear correlations (significance level, $\alpha = 0.05$), respectively, and shaded areas represent the 95% confidence interval of the predictions. In panel a, the niche specialization was Box-Cox transformed, and then was rescaled to the range [0, 1].

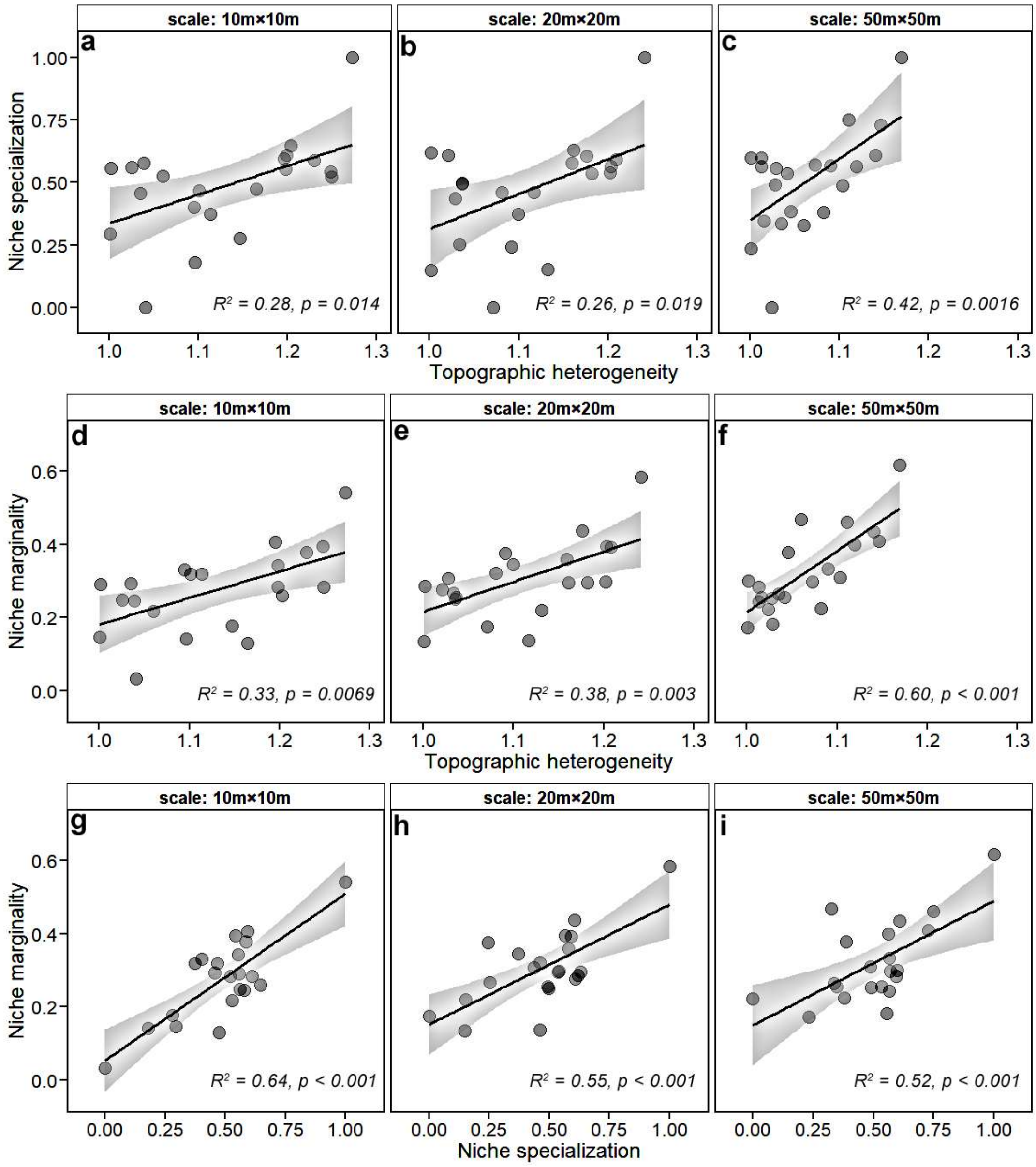


**Supplementary Figure S4**. **Relationships between topographic heterogeneity, community-level niche specialization and niche marginality across grain sizes (a, d, and g: 10 m × 10 m; b, e, and h: 20 m × 20 m; c, f, and i: 50 m × 50 m).** The niche specialization was transformed into normality using a Box-Cox transformation, and then was rescaled to the range in [0, 1] with the min-max normalization. Topographic heterogeneity was quantified as surface: Planimetric area ratio.

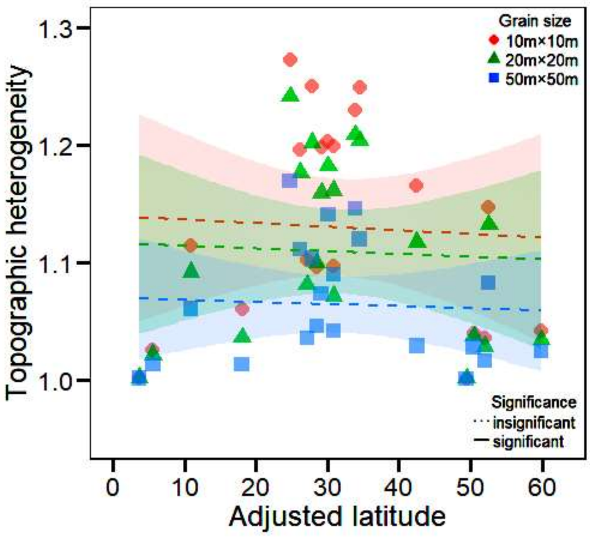


**Supplementary Figure S5**. **The linear relationship between topographic heterogeneity (quantified by the surface to planimetric area ratio) and adjusted latitude across grain sizes (10 m × 10 m, 20 m × 20 m and 50 m × 50 m).** Dashed lines indicate insignificant linear correlations (significance level, $\alpha$ = 0.05), and different colours of points and lines represent grain sizes.

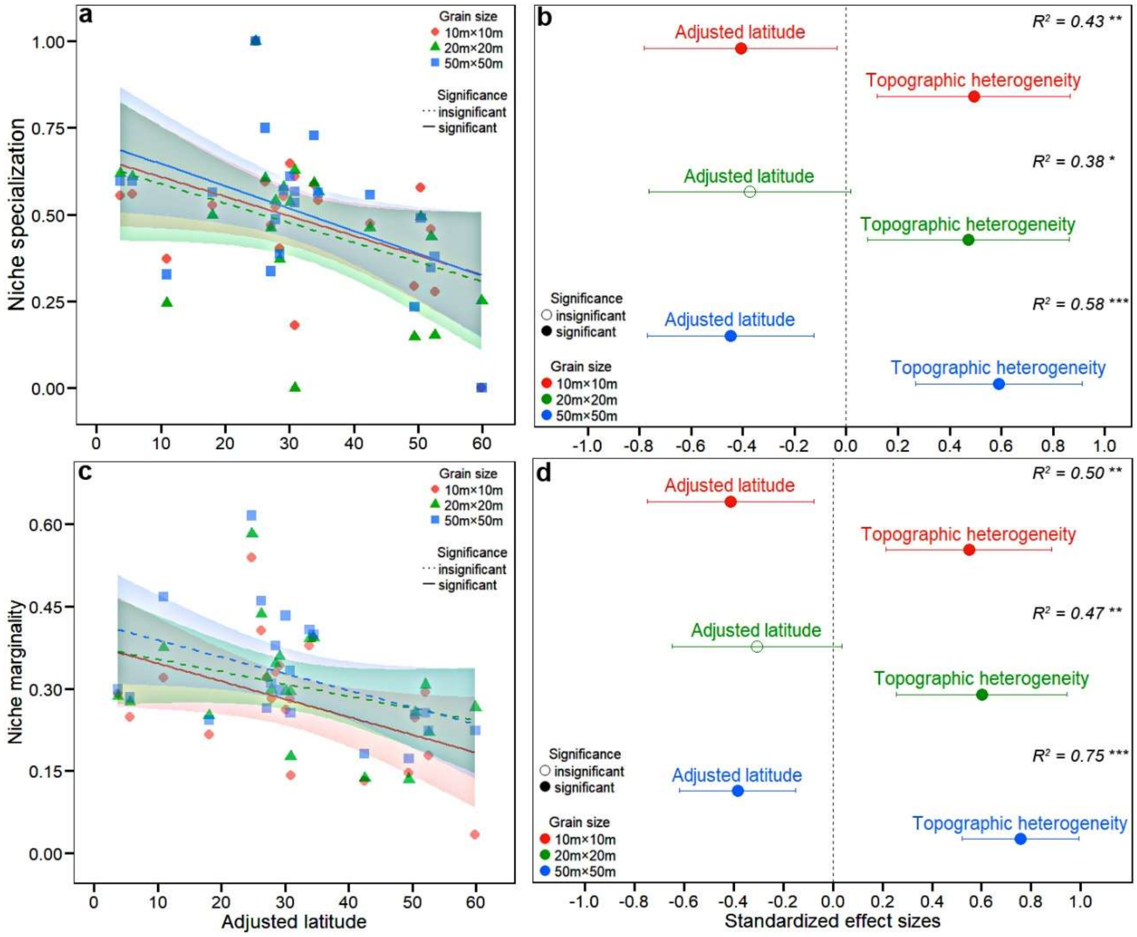


**Supplementary Figure S6. The relationships of community-level niche specialization (a and b) and marginality (c and d) with adjusted latitude and local topographic heterogeneity across grain sizes.** Community-level niche specialization was Box-Cox transformed and was subsequently scaled to the range [0, 1] for comparison across grain sizes. In each panel, different colours of points and lines represent grain sizes. In panels a and c, solid and dashed lines indicate significant and non-significant linear correlations ($\alpha = 0.05$), respectively, and shaded areas represent the 95% confidence intervals of the predictions (electronic supplementary material, table S4). In panels b and d, points represent the standardized effect sizes of explanatory variables that are significantly (solid circles) and non-significantly (open circles) different from zero, respectively. The significance level of the total $R^2$ are $\alpha < 0.001$, '***'; α < 0.01, '**'; α < 0.05 '*'.
